# Experimental infection of mink with SARS-COV-2 Omicron (BA.1) variant leads to symptomatic disease with lung pathology and transmission

**DOI:** 10.1101/2022.02.16.480524

**Authors:** Jenni Virtanen, Kirsi Aaltonen, Kristel Kegler, Vinaya Venkat, Thanakorn Niamsap, Lauri Kareinen, Rasmus Malmgren, Olga Kivelä, Nina Atanasova, Pamela Österlund, Teemu Smura, Antti Sukura, Tomas Strandin, Lara Dutra, Olli Vapalahti, Heli Nordgren, Ravi Kant, Tarja Sironen

**Author notes:** These authors contributed equally to this article (Jenni Virtanen, Kirsi Aaltonen). Address for Correspondence: Jenni Virtanen, Department of Veterinary Biosciences, Faculty of Veterinary Medicine, P.O. Box 66, and Department of Virology, Faculty of Medicine, P.O. Box 21, 00014 University of Helsinki, Helsinki, Finland.

## Abstract

We report an experimental infection of American mink with SARS-CoV-2 Omicron variant and show that minks remain virus RNA positive for days, develop clinical signs and histopathological changes, and transmit the virus to uninfected recipients warranting further studies and preparedness.

## Text

SARS-CoV-2 has been detected in farmed and feral American mink in multiple countries with evidence of extensive environmental contamination and human-to-mink and mink-to-human transmission *(1-5*). This has led to strict measures in mink farms and mink farming countries to prevent the spread of the disease. In late 2021, a new SARS-CoV-2 variant (Omicron), characterized by possibly milder symptoms and more efficient human-to-human transmission, was detected, but its infectivity and spread in American mink in unknown *(6, 7*).

We tested the response of the American mink to the Omicron variant by infecting three male mink intranasally with 4 × 10^5^ PFU of the virus (see Appendix for methods). Infected minks were followed for seven days, sampled daily for saliva, and subjected to histopathological evaluation of upper and lower respiratory tracts on the last day of follow-up.

All experimentally infected mink showed mild to moderate signs of illness including lethargy, anorexia, diarrhea, nasal and lacrimal discharge, and sneezing. Consistent with earlier experiments with other variants (*8, 9*), saliva samples tested PCR positive 1-day post-infection (dpi) and remained that way throughout the follow-up (Table and Appendix). Even though some of the clinical signs may be due to other factors such as stress from the change of environment, consistency of symptoms to studies with other variants combined with PCR results, demonstrate that the Omicron variant also causes a symptomatic infection in mink.

**Table:**
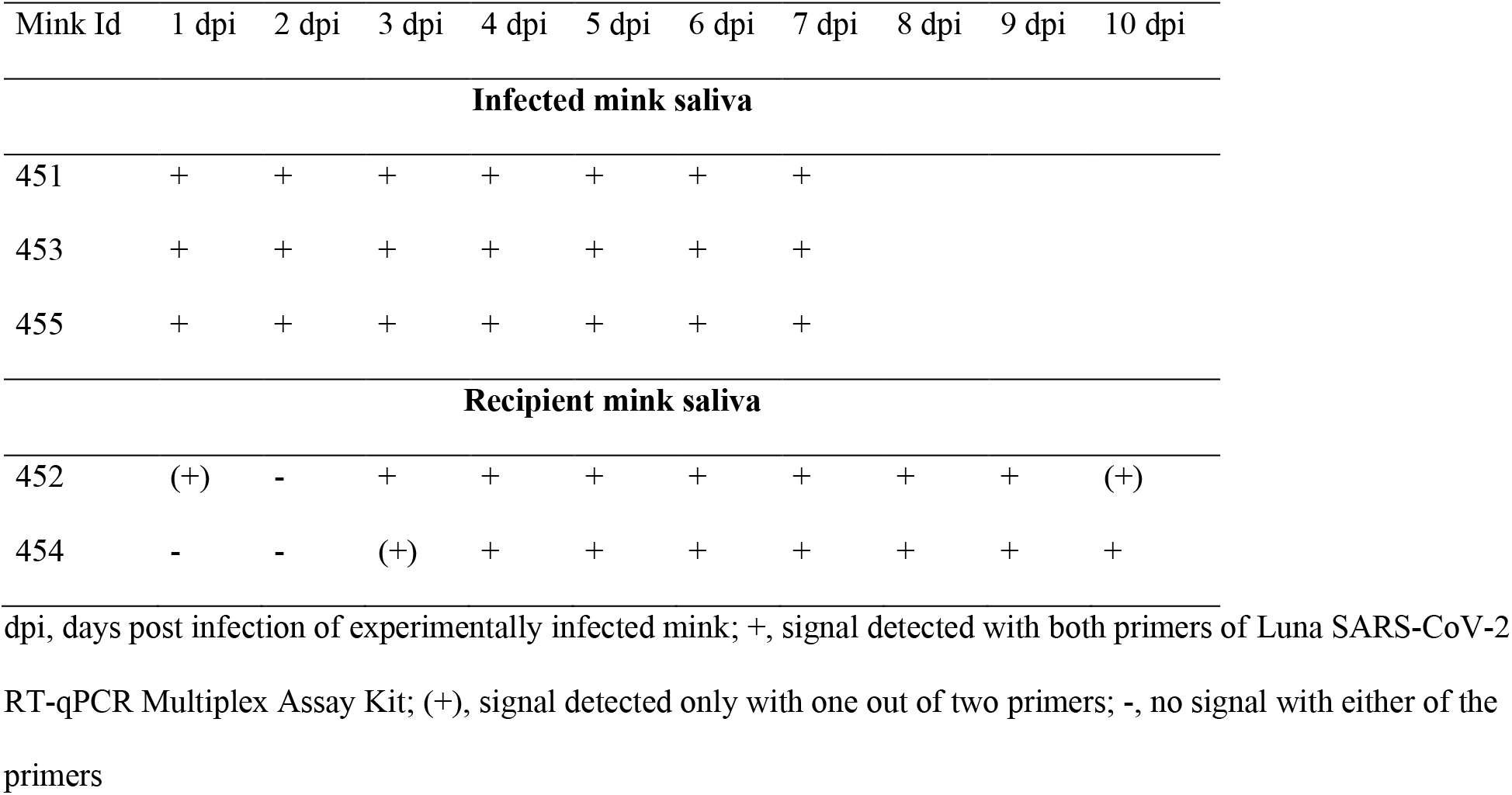
PCR results from saliva of three experimentally infected mink and two uninfected recipient mink

| Mink Id | 1 dpi | 2 dpi | 3 dpi | 4 dpi | 5 dpi | 6 dpi | 7 dpi | 8 dpi | 9 dpi | 10 dpi |
| --- | --- | --- | --- | --- | --- | --- | --- | --- | --- | --- |
| **Infected mink saliva** |  |  |  |  |  |  |  |  |  |  |
| 451 | + | + | + | + | + | + | + |  |  |  |
| 453 | + | + | + | + | + | + | + |  |  |  |
| 455 | + | + | + | + | + | + | + |  |  |  |
| **Recipient mink saliva** |  |  |  |  |  |  |  |  |  |  |
| 452 | (+) | - | + | + | + | + | + | + | + | (+) |
| 454 | - | - | (+) | + | + | + | + | + | + | + |

dpi, days post infection of experimentally infected mink; +, signal detected with both primers of Luna SARS-CoV-2 RT-qPCR Multiplex Assay Kit; (+), signal detected only with one out of two primers; -, no signal with either of the primers

To study if mink can transmit the virus, two minks were used as uninfected recipients in separate cages located 10-20 cm away from the cages of the infected mink and followed for ten days. Both recipients developed similar symptoms to the experimentally infected mink and were consistently PCR positive from day three onwards (Table), indicating mink-to-mink transmission. Even though there is currently no evidence of mink-to-human transmission of the Omicron variant, it seems likely based on our results and the information from other variants.

Gross findings in the nasal cavity and lungs were subtle in both experimentally infected and recipient mink consisting of hyperemia of respiratory mucosa with small amounts of viscous exudate and non-collapsed, dark-red, and wet pulmonary lobes. All mink showed consistent histopathological changes in the upper and lower respiratory tracts. Multifocal degeneration and loss of respiratory epithelium with variable mucosal and submucosal neutrophilic infiltration was observed in the nose. The lumen contained sloughed epithelial cells, mucinous material, and degenerated neutrophils (Figure A, C). Viral nucleoprotein was widely distributed beyond intact cells, within sloughed cells, and mucosal respiratory epithelium (Figure B, D). The olfactory epithelium was inconsistent and mildly affected with only focal viral antigen detection. Unlike in some experimental infections reported in rodents, clear pathology was observed in the lungs (*10*). In two inoculated and both recipient mink, pulmonary lesions (Figure E, G) were associated with viral antigen expression (Figure F, H) and characterized by multifocal to coalescing alveolar damage with degeneration and/or necrosis of alveolar septa, infrequent hyalin membrane formation, and variable proliferation of type II pneumocytes (Figure I, J). Alveolar spaces contained macrophages, sloughed cells, edema, and hemorrhage. Bronchiolar epithelial degeneration and hyperplasia were variably present (Figure K), and the lumen filled with few sloughed cells and neutrophils. Bronchi were lined by hyperplastic epithelium with increased numbers of goblet cells. Other consistent findings were vasculitis (Figure L), perivasculitis, perivascular, and peri-bronchial edema. One inoculated mink had markedly thickened alveolar septa by mononuclear cells, marked proliferation of type II pneumocytes, intra-alveolar macrophages, few syncytial cells, bronchi and bronchiolar epithelial cell hyperplasia, vasculitis, and perivasculitis. Viral antigen could not be detected in this mink. Strikingly, all evaluated mink lacked viral antigen in the epithelium of bronchi and bronchioles.

**Figure.**
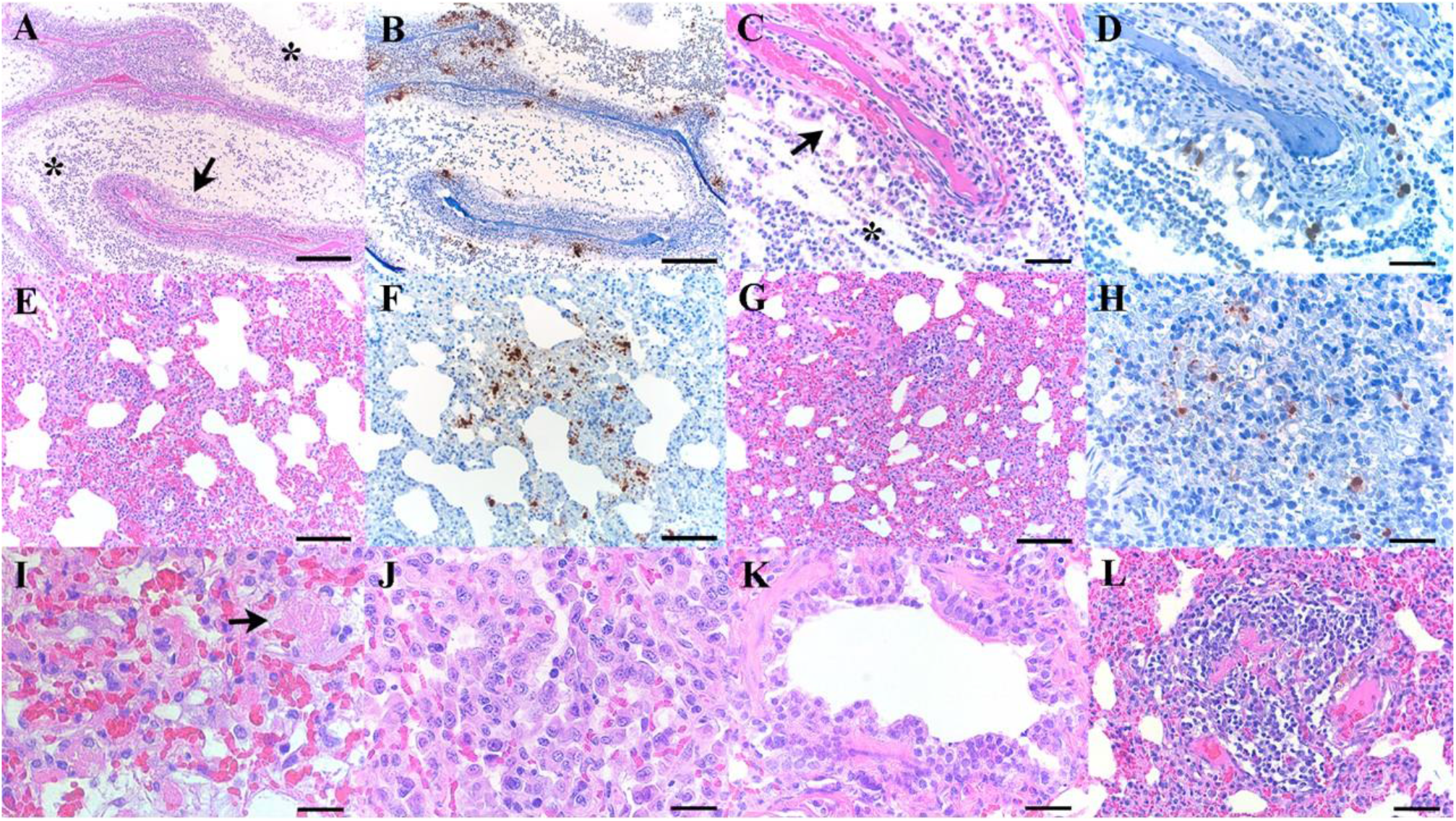
Histopathological changes and SARS-CoV-2 expression in the upper and lower respiratory tracts in experimentally infected mink with Omicron variant at 7 dpi and recipient mink after 10 days of follow up. A) Respiratory segment of the nose from an intranasally infected mink showing luminal accumulation of exudate (*) and degeneration of mucosal epithelium (arrow, bar 500μm. B) Viral antigen is widely detected within nasal lumen and respiratory epithelium (bar 500μm). C) Respiratory epithelium from a recipient mink depicting marked degeneration and loss (arrow), intraluminal accumulation of sloughed cells and neutrophils (*), and D) intraepithelial viral expression (C, D bar 50 μm). E-H) Lungs from intranasally infected (E, F) and recipient mink (G-H) showing alveolar damage with intralesional presence of viral nucleoprotein (E-G bar 200μm, H bar 50 μm). I, J) Marked degeneration and necrosis of alveolar septa and focal hyalin membrane (arrow, I), and prominent proliferation of type II pneumocytes (J) in an intranasally infected mink (I, J bar 25μm). K, L) Recipient mink showing bronchiolar epithelial degeneration and hyperplasia (K), and vasculitis (L) with complete destruction of blood vessel wall and mononuclear cell infiltration (K bar 50μm, L bar 100μm). Haematoxylin and esosin (HE) stain and immunohistochemistry, haematoxylin counterstain.

The Omicron variant is different from other variants due to its more efficient spread, primarily attributable to immune escape and likely milder symptoms in humans (*6, 7*). This makes it more difficult to prevent virus introduction into the mink farms through asymptomatic humans, creating a more significant risk for the formation of virus reservoirs among farmed or feral mink. This study shows that mink can be infected by Omicron and importantly, efficiently transmit the virus to other mink. Despite the reports of lower virulence of Omicron, mink develop clinical disease and nasal and pulmonary microscopic lesions closely resemble infection with previously reported variants in mink and humans. A better understanding of the clinical symptoms helps detect the virus among mink earlier. Further studies are needed to determine the risk of transmission to humans, emergence of mink-specific mutations, the pathogenesis of pulmonary involvement, and prepare for this easily transmitted variant among farmed and feral mink.

## Acknowledgements

We thank animal Jari Elemo and Mari Elemo and other animal care takers for handling the animals and assessing their health and Esa Pohjalainen, Sanna Mäki, Tiina Sihvonen, Johanna Rintamäki, Hanna Valtonen, Marika Skön, Larissa Laine, and Elina Väisänen for technical assistance. We thank Kati Kuipers, Anne Kujanpää, Laura Vähälä, and the Finnish Centre for Laboratory Animal Pathology (FCLAP) for expert technical help as well as Johanna Korpela and Jussi Peura from Finnish Fur Breeders Association and Jan Segervall and Maarit Mohaibes from the Kannus Research Farm Luova Ltd. for providing the animals. We also thank E3 Excellence in Pandemic Response and Enterprise Solutions co-innovation project and all its parties.

This study was supported by the Academy of Finland (grant number 336490, 339510), VEO - European Union’s Horizon 2020 (grant number 874735), Business Finland E3 (4917/31/2021), Finnish Institute for Health, and Welfare and the Jane and Aatos Erkko Foundation.

