## Appendix for "Experimental infection of mink with SARS-COV-2 Omicron (BA.1) variant leads to symptomatic disease with lung pathology and transmission"

### Materials and methods

**Virus stock**

Omicron variant (original patient sample: hCoV-19/Finland/THL-202126660/2021, EPI_ISL_8768822 (Gisaid)) of SARS-CoV-2 (10^6 PFU/ml) was acquired from the Finnish institute of Health and Welfare. We ensured that the viral genome in the stock (OM393712) remained unchanged, including the furin cleavage site that mutates particularly rapidly in cell culture for Omicron, with a protocol described in (*1*). It has been reported that a few key mutations (e.g. the N501Y mutation) present in alpha, beta, gamma, and omicron variants improve the affinity of the SARS-CoV-2 spike S1 protein to mouse ACE-2 and increase infectivity of SARS-CoV-2 in standard BALB/c mice (*1*).

**Animals, infection, and euthanasia**

A total of 5 male American mink (Neovison vison) were transported to the University of Helsinki biosafety level-3 (BSL-3) facility and acclimatized to custom made cages with nest boxes for three days with ad libitum water and food. For the experimental infection, mink were anesthetized with 30 µl of Ketaminol (100 mg/ml, Intervet) and Domitor (1 mg/ml, Orion Pharma) via i.m. administration and three were inoculated intranasally with 200 µL of virus stock into both nostrils and the remaining two with phosphate buffered saline (PBS). The animals were held in an upright position for a few seconds to allow the liquid to flush downwards in the nasal cavity. Revertor (5 mg/ml, Scanvet) was given (15ul) as an α2-adrenergic antagonist for reversing clinical effects of sedation. All minks were monitored daily for signs of illness (changes in posture or behavior, rough coat, apathy, ataxia, runny nose, diarrhea etc.). At the end of the experiment, the minks were anesthetized with Ketaminol and Domitor (40 µl), sampled for blood, and euthanized in a CO2 chamber. Experimental procedures were approved by the Animal Experimental Board of Finland (license number ESAVI/33259).

**Sample collection**

Saliva samples (oral swabs) were collected in triplicates before the infection, and everyday post infection with foam swabs (Virocult, MWE) into 200 µl PBS. Animals were necropsied immediately after euthanasia. Representative fresh samples from all lung lobes were collected and frozen for virological examinations. Lungs were then inflated with 10% neutral buffered formalin and fixed for 48 h. For histopathology, each lung lobe (right cranial, left cranial, right medial, right caudal and left caudal) were trimmed into three consecutive cross sections of approximately 0.5 cm thick starting at the lobe hilum and moving toward the posterior of the lobe along the main branching bronchus. The head was separated from the carcass by disarticulation of the atlanto-occipital joint, the skull was removed exposing the brain and the entire was placed in 10% neutral buffered formalin. After 48 h fixation, the brain was removed and the head sawn longitudinally in the midline using a diamond saw. Slices of 0.3 cm were prepared from the nasal cavity including vestibular, respiratory and olfactory segments, and gently decalcified for 4 days in 14% EDTA solution (Tritrimex ® III, Nederland).

##### **Histology and Immunohistochemistry**

Trimmed lung lobes and decalcified nasal cavity sections were routinely processed, paraffin-wax embedded, cut at 3–5 µm and stained with haematoxylin–eosin (HE) or subjected to immunohistochemistry (IHC) for the detection of SARS-CoV-2 antigen. Briefly for IHC, slides were deparaffinized and incubated for 20 min at 99°C in 10 mM citrate buffer (pH 6) for heat-induced antigen retrieval. Endogenous peroxidase was performed by immersion in 3% hydrogen peroxide for 10 min. After incubation with 10% bovine serum albumin in phosphate-buffered saline, sections were incubated for 60 minutes at room temperature (RT) with rabbit polyclonal anti- SARS-CoV-2 nucleoprotein antibody diluted 1:3000 in animal-free blocker and diluent solution (R.T.U. Animal-Free Blocker and Diluent; Vector Laboratories, Burlingame, Ca, USA). Polymer-linked to HRP (BrightVision + Poly-HRP kit; ImmunoLogic, Duiven, Netherlands) was used as secondary antibody incubating in a humid chamber for 30 min at RT. The reaction was visualized with right DAB Substrate kit (ImmunoLogic, Duiven, Netherlands) and slight counterstain with Harris hematoxylin. As negative control, the primary antibody was substituted with rabbit IgG Isotype control (1:1000).

**PCR and sequencing**

RNA was extracted from saliva samples collected in PBS with Viral RNA Minikit (Qiagen) according to manufacturer’s instructions. PCR tests were performed with Luna SARS-CoV-2 RT-qPCR Multiplex Assay Kit (NEB). Samples that gave signal with both primers were considered positive (+) and samples that only gave signal with one primer weak positive ((+)).

**Appendix Table.** Ct values from saliva samples in PCR with Luna SARS-CoV-2 RT-qPCR Multiplex Assay Kit (2019-nCoV_N1/2019-nCoV_N2)

|  | Infected mink | | | Recipient mink | |
| --- | --- | --- | --- | --- | --- |
| dpi | 451 | 453 | 455 | 452 | 454 |
| 1 | 28.15/28.27 | 23.27/23.85 | 25.58/25.66 | no ct/36.67 | no ct/no ct |
| 2 | 28.23/28.55 | 34.63/34.17 | 29.66/30.16 | no ct/no ct | no ct/no ct |
| 3 | 14.15/13.86 | 14.9/14.61 | 15.85/15.71 | 33.36/33.38 | 38.68/no ct |
| 4 | 25.3/26.27 | 21.19/21.99 | 25.66/26.72 | 28.71/30.06 | 31.13/32.27 |
| 5 | 32.44/33.22 | 25.87/26.00 | 23.71/23.83 | 34.13/34.12 | 26.46/26.59 |
| 6 | 27.49/28.01 | 29.79/30.77 | 32.17/33.97 | 30.19/30.75 | 33.57/34.16 |
| 7 | 25.03/26.91 | 27.07/28.78 | 27.82/29.80 | 30.91/33.51 | 27.25/29.12 |
| 8 | N/A | N/A | N/A | 28.07/28.29 | 31.81/32.01 |
| 9 | N/A | N/A | N/A | 30.69/32.56 | 34.7/36.36 |
| 10 | N/A | N/A | N/A | 37.41/no ct | 32.79/34.05 |
